# DNA methylation in the upstream CpG island of the GPER locus and its relationship with GPER expression in colon cancer cell lines

**DOI:** 10.1101/2020.07.03.187351

**Authors:** Uttariya Pal, Sujasha Ghosh, Anil Mukund Limaye

**Affiliations:** Department of Biosciences and Bioengineering, Indian Institute of Technology Guwahati, Guwahati 781039, Assam, India

**Author notes:** Corresponding author Dr. Anil Mukund Limaye, Department of Biosciences and Bioengineering, Indian Institute of Technology Guwahati, Guwahati 781039, Assam, India.

**Keywords:** GPER, methylation, bisulfite sequencing, colon cancer

## Abstract

The seven-transmembrane G-protein coupled estrogen receptor (GPER) relays short-term non-genomic responses in target cells and tissues. It is a proposed tumor suppressor, which frequently undergoes down-modulation in primary tumors of the breast, ovary, and endometrium. A study by Liu and co-workers reported the loss of GPER expression in colorectal cancer and attributed it to DNA methylation-dependent silencing. The present study is based on the hypothesis that GPER expression is inversely correlated with methylation in the upstream CpG island (upCpGi) in the GPER locus. Methylation in the upCpGi was analysed by bisulfite sequencing and correlated with GPER expression in a panel of colon cancer cell lines The bisulfite sequencing results show the presence of a differentially methylated region (DMR) comprising of the downstream eight CpGs of the upCpGi. Methylation in the DMR correlated inversely with GPER expression. We compared two cell lines, namely SW620 and COLO-320DM, in terms of their viability in response to varying concentrations of G1, a GPER specific agonist, which is known to induce cell cycle arrest and apoptosis in colon cancer cell lines. SW-620 cells, which had the least methylated DMR and the highest level of GPER expression, showed significant loss of viability with 1 μM G1. On the other hand, COLO-320DM, which had the most methylated DMR and the lowest level of GPER expression, did not show a significant response to 1 μM G1. At 5 μM G1, SW620 cells showed a greater reduction in viability than COLO-320DM cells. Our study demonstrates the inverse correlation between DNA methylation in the DMR and GPER expression. GPER is a non-canonical form of estrogen receptor, and estrogen is believed to exert its oncoprotective effect in the colon via GPER. DNA methylation-dependent silencing of GPER may, at least in part, the underlying reason behind the loss of estrogen’s oncoprotective effect in the colon. Future studies should explore the utility of DNA methylation in the upCpGi, particularly the DMR, in diagnosis or prognosis.

## INTRODUCTION

Colorectal cancer is the second most diagnosed cancer in both men and women. It is also the second most leading cause of cancer-related deaths (1). The incidence of colorectal cancer in women is significantly lower compared to men (2). Women receiving hormone replacement therapy have a significantly lower risk of colorectal cancer (3,4). Premenopausal women diagnosed with colorectal cancer have better survival than age-matched men (5). These findings posit estrogen as an important etiological factor in the colorectal tissue and indicate its oncoprotective action.

Estrogen exerts its effects on target cells and tissues via genomic and non-genomic pathways. The genomic effects of estrogen are mediated by the canonical estrogen receptors, namely ERα and ERβ. These are ligand-dependent transcription factors encoded by ESR1 and ESR2 genes, respectively (6). The non-genomic effects of estrogen are mediated by membrane-tethered canonical estrogen receptors (7–10), ERα36 (11,12), a splice variant of ERα, or non-canonical estrogen receptors such as the G-protein coupled estrogen receptor (GPER) (13). Low ERα expression in the colon makes it an unlikely mediator of estrogen actions in this tissue (14). ERβ is abundantly expressed in the normal colon (15). Its expression is reduced in colorectal cancer, and its loss is associated with higher Dukes’ stages and poor prognosis (16–19). ERβ knockout mice develop colitis-associated neoplasia (20). Thus, ERβ may be a crucial mediator of estrogen’s oncoprotective effects in the colon, and a target for colorectal cancer prevention (21).

The G protein-coupled estrogen receptor (GPER, formerly known as GPR30) is the most recent entry to the list of membrane-associated ERs (mER). Upon estrogen binding, it produces short-term non-genomic effects, such as increased cAMP, increased cytoplasmic calcium, and activation of PI3K and MAPK. GPER has been studied in almost all physiological systems, such as immune, reproductive, cardiovascular, neuroendocrine, urinary and musculoskeletal(22,23). It is an emerging prognostic marker and potential therapeutic target in endocrine cancers(24–29).

Recent studies provided insights into the role and relevance of GPER in the physiology and pathology of the colonic environment. They projected it as a significant mediator of estrogen actions in the colon (30). The importance of GPER expression, or lack of it, in the etiology of colorectal cancer and its progression is ambiguous. Liu and co-workers reported reduced GPER expression in colorectal cancer samples compared to normal counterparts. Reduced GPER expression was associated with poor prognosis, suggesting that GPER is a tumor suppressor (31). On the other hand, hypoxia induces GPER expression. Bustos and co-workers found evidences that support the pro-tumorigenic role for GPER, especially in the face of reduced ERβ expression in the hypoxic tumor microenvironment (32). Despite the contradiction, the association between altered GPER expression and colorectal cancer is apparent.

Recent studies, including those reported from our laboratory, show that epigenetic silencing is, at least in part, responsible for the loss of GPER expression in breast and colorectal cancer (31,33–36). In a study on breast cancer cell lines as models, we found that GPER expression had an inverse relationship with methylation in the upstream CpG island (upCpGi) in the GPER locus. We also described a differentially methylated region (DMR) comprising of terminal eight CpG dinucleotides in the upCpGi, which was differentially methylated in two breast cancer cell lines with contrasting GPER expression levels (35). The methylation status in the DMR has not been examined in colorectal cancer. In the present study, we have examined GPER expression and upstream CpG island methylation in a panel of colon cancer cell lines.

## MATERIALS AND METHODS

### Plasticwares, chemicals, and reagents

Cell culture plasticwares were purchased from Eppendorf (Hamburg, Germany). Dulbecco’s modified eagle’s medium (DMEM) and Roswell Park Memorial Institute (RPMI) 1640 with phenol red were purchased from HiMedia (Mumbai, India). Fetal bovine serum (FBS) was from Invitrogen Corporation (Grand Island, NY, USA). Trypsin-EDTA, antibiotics, and Dulbecco’s phosphate-buffered saline (DPBS) were purchased from HiMedia (Mumbai, India). EmeraldAmp RR320B MAX PCR Master Mix was from Takara Bio Incorporation (New Delhi, India). The GPER-specific agonist, G1, was purchased from Cayman Chemical (Cat. No. CAS 881639-98-1) (MI, USA). All other chemicals, salts, and buffers were from Merck and SRL (Mumbai, India).

### Cell culture

Colon cancer cell lines HT-29, HCT-15, HCT-116, SW-480, SW-620, COLO-205, COLO-320DM,and breast cancer cell line MCF-7 were obtained from the National Centre for Cell Science (NCCS,Pune, India). They were routinely cultured and expanded under standard conditions of 37°C and 5% CO_2_ in phenol red-containing DMEM for HT-29 and MCF-7, and RPMI 1640 for the remaining cell lines. Media were supplemented with 10% heat-inactivated FBS, 100 units/mL penicillin, and 100 μg/mL streptomycin.

### Total RNA, Protein isolation and cDNA synthesis

Total RNA and protein were isolated from cell lines as described previously (35). 2 µg of total RNA was reverse transcribed using High Capacity cDNA Reverse Transcription kit (Invitrogen, USA) as per manufacturer’s instructions.

### RT-PCR and western blotting

The methodologies for RT-PCR and western blotting, including the primer and antibody details are as described earlier (35,37). RPL35a and Histone H3 were used as internal controls in RT-PCR and western blotting analyses, respectively.

### Bisulfite sequencing

Genomic DNA (gDNA) from the cell lines was extracted using PureLink Quick Gel Extraction Kit (Invitrogen, CA, USA) and subjected to bisulfite conversion using the EpiJET Bisulfite Conversion kit (Thermo Fisher Scientific, USA) as per manufacturer’s instructions. The bisulfite-converted gDNA was used as a template to generate upCpGi containing amplicons using specific primers reported earlier (35). The amplified products were eluted from agarose gels and ligated into the pMD20 vector (Takara Bio, India) as per the manufacturer’s instructions. The ligated products were then transformed into *E.coli* Top10 competent cells, and recombinant colonies screened using ampicillin resistance as the selectable marker. Plasmids were isolated using GSure Plasmid Kit (GCC Biotech, India), and the inserts of 10 independent recombinant plasmids per cell line were sequenced. Methylated and unmethylated CpG sites were identified and represented as lollipop plots. The proportion of CpGs methylated in the upCpGi was determined for each cell line. The sequencing was done by Eurofins Scientific (Bangalore, India).

### G1 Treatment and determination of cell viability

Colon cancer cells were seeded (4000-5000 cells/well) in 96 well plates. After 48 h, the monolayer was washed with DPBS. The cells were then processed directly for MTT assay to determine baseline viability or treated with indicated concentrations of G1 or ethanol (vehicle control) for 120 h before MTT assay. In case of the latter, media containing G1 or ethanol were replenished after every 48 h. MTT assays were performed as described previously (38). For each cell line, the change in viability due to treatment by ethanol or G1 was determined by subtracting the baseline viability. The change in viability of ethanol treated cells were assigned the value of 100, and those of G1 treated cells were expressed relative to control. COLO-205 cells, which are semi-adherent cells, were not amenable to MTT assays.

### Statistical analysis

Data generated from MTT assays were analyzed using one-way ANOVA followed by Tukey’s HSD using the R statistical package. Methylation in the upCpGi of GPER locus and comparison of percent methylation across all cell lines were performed as described earlier (39). In all the tests, p<0.05 was considered statistically significant.

## RESULTS

### GPER expression in colon cancer cell lines

There are three variants of GPER mRNA, namely NM_001505.2 (GPERv2), NM_001039966.1 (GPERv3) and NM_001098201.1 (GPERv4). They have exactly the same open reading frame and 3’UTR, but have different exon-intron organization and 5’UTR. Using variant-specific primer combinations described earlier (35), we studied the expression of GPER mRNA in a panel of colon cancer cell lines. MCF-7 cell line which expresses the three variants (35), was used as a reference. The colon cancer cell lines expressed varying degrees of GPER mRNA variants (Fig 1A). In SW-480 and SW-620 cells, the GPERv2 and GPERv3 were the dominant variants. GPERv3 was the major variant in HT-29 cells with very low and undetectable levels of GPERv4 and GPERv2, respectively. HCT-15 showed almost no expression of GPER mRNA. Other cell lines had a moderate expression of GPER mRNA variants (Fig 1A). We examined GPER protein expression in these cell lines. The protein levels, by and large, were consistent with the levels of GPER mRNA (Fig 1B). HT-29 was an exception. Despite appreciable level of GPERv3, negligible protein could be detected (Fig 1A and 1B).

**Figure 1.**
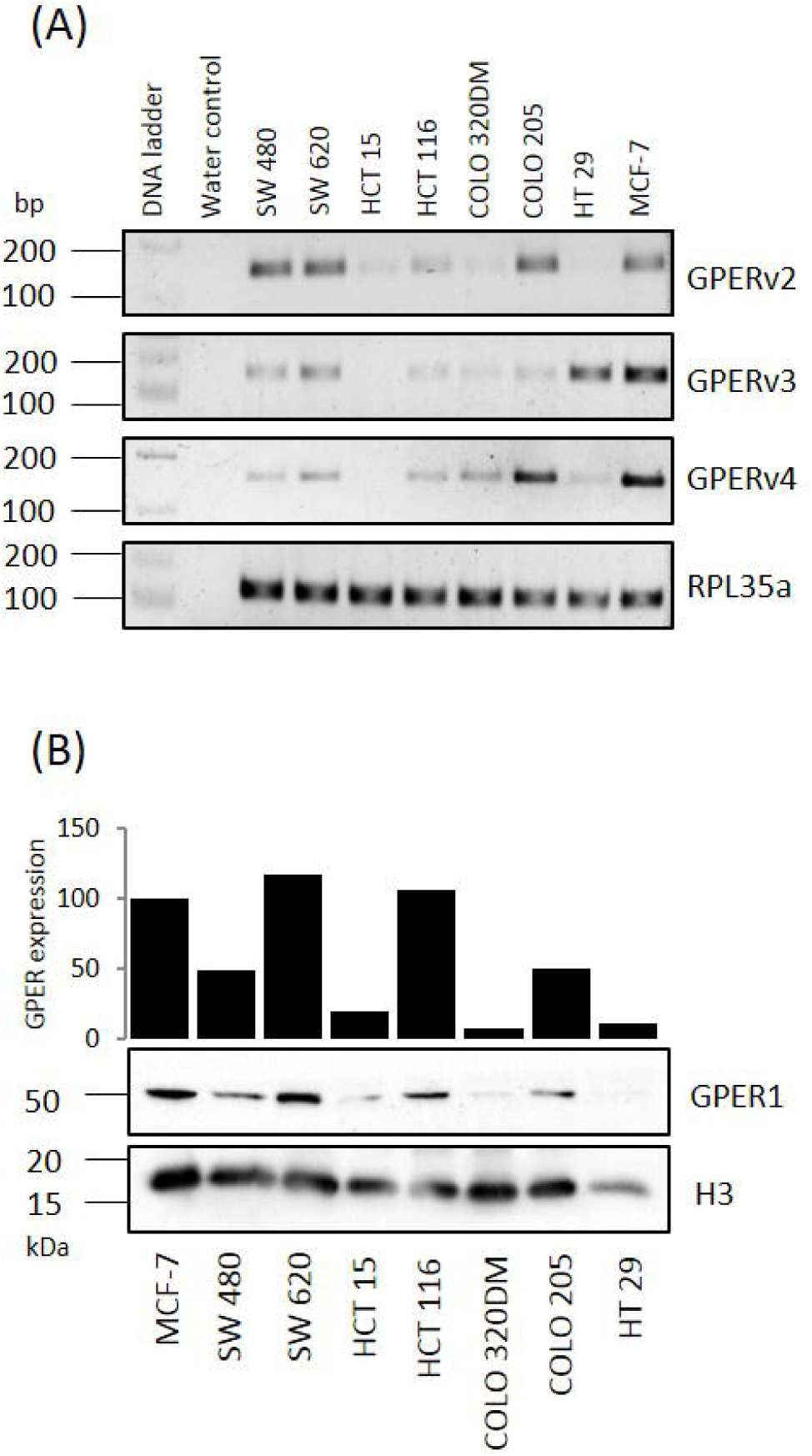
GPER expression in colon cancer cell lines. Total RNA and protein were isolated from the indicated cell lines. GPER expression was determined by RT-PCR (A) using primers and PCR conditions described previously (35), and western blotting (B). The protocol for western blotting was as described earlier (35,37).

### CpG methylation in the upCpGi

Using bisulfite sequencing approach as described earlier (35), we analysed DNA methylation in the upCpGi, which harbors 32 CpG sites. Figure 2A shows the methylation pattern deduced from bisulfite sequencing data in the form of lollipop models. The upCpGi in COLO-320DM was extensively methylated, which was in sharp contrast to the observation in SW-620 cells. The remaining cell lines showed varying levels of methylation, but not as extensive as that in COLO-320DM. Interestingly, much of the methylation was observed in the downstream terminal eight CpGs in the upCpGi (referred to as the DMR). This was similar to our previous observation in MCF-7 and MDA-MB-231 breast cancer cell lines (35). The proportion of methylated CpGs in the DMR for each of the cell lines is indicated in parenthesis in Fig 2A. For each pair of cell lines, the difference in the proportion of methylated CpGs was tested for statistical significance. The adjusted p-values are indicated in Fig 2B. The level of GPER protein significantly correlated inversely with the proportion of methylated CpGs in the DMR (Fig 2C, Spearman’s rho = −0.78, p = 0.048).

**Figure 2.**
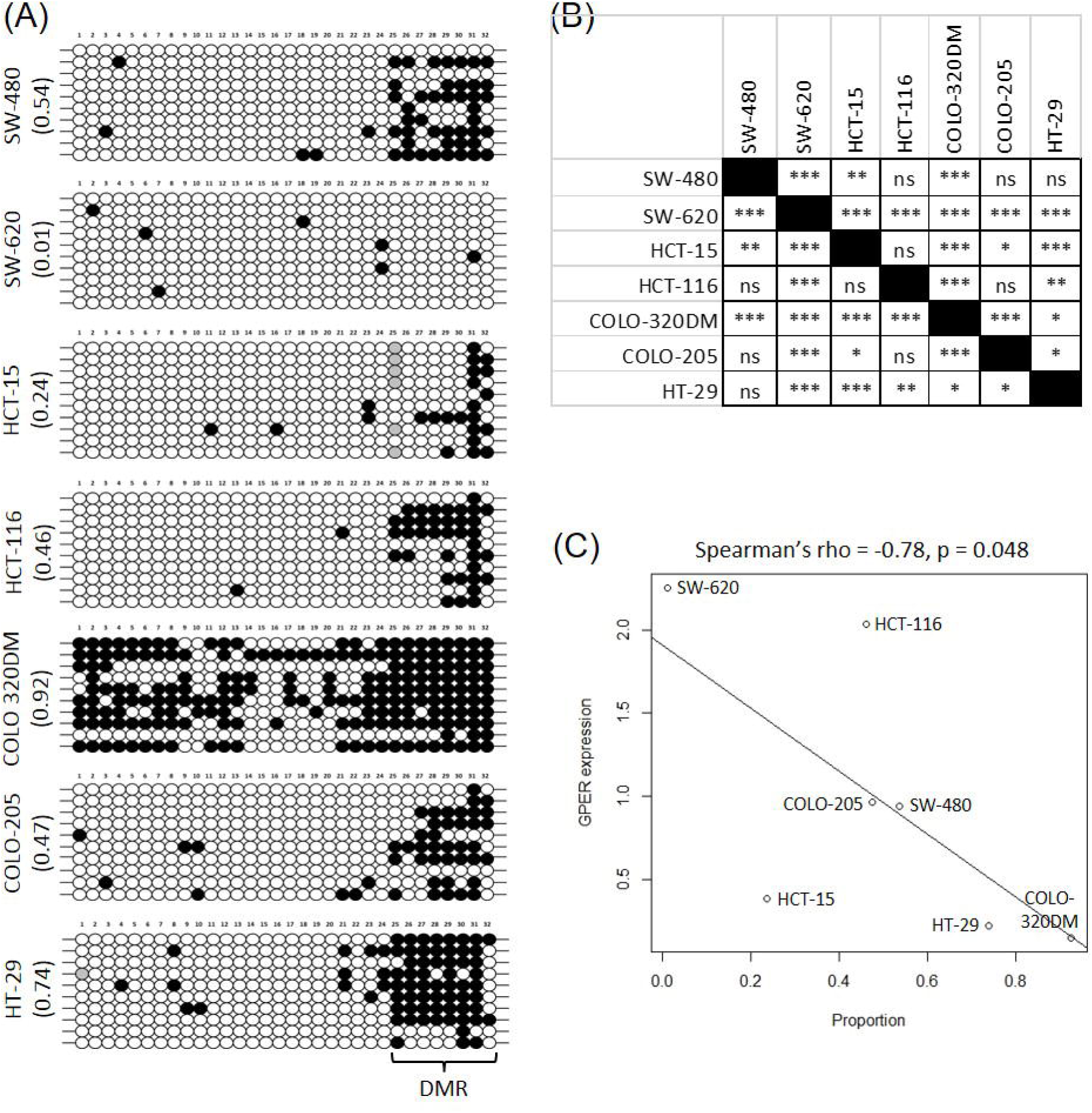
upCpGi methylation in the GPER locus. Genomic DNA isolated from the indicated colon cancer cell lines were bisulfite converted and used for PCR reaction with primers described earlier to amplify the upCpGi. The PCR amplified products were cloned in pMD20. Inserts from 10 independent TA clones per cell line were sequenced and analyzed for methylated and unmethylated CpG sites. There are 32 CpG sites in the upCpGi. A. Lollipop models depicting the status of each CpG site in the indicated cell lines. Filled circles represent methylated CpGs, and Open circles represent unmethylated CpGs. Grey circles represent the absence of a CpG site. Since there are 8 CpGs in the DMR, a total of 80 CpGs were sampled for each cell line (n=10 clones for each sample). The numbers in parentheses denote the proportion of methylated CpGs sampled in the DMR. B. Adjusted p-values for the test of significant difference in proportion of CpGs methylated in the DMR. *p < 0.05, **p < 0.01, ***p<0.001, ns – not significant. C. Relationship between the proportion of CpGs methylated in the DMR and GPER expression in colon cancer cell lines.

### Response to GPER stimulation

G1, a specific GPER agonist, induces cell cycle arrest and apoptosis in colon cancer cell lines. We tested whether the response of the cell lines under study to G1 treatment was commensurate with the levels of GPER protein expression. We designed an MTT assay to examine the change in viability of cells following 120 h of treatment with varying concentrations of G1 after due correction of the zero-hour baseline viability. Up to 500 nM G1, none of the cell lines had reduced viability (Fig 3), except HCT15. HCT15 had significantly reduced viability with 500 nM G1 (Fig 3C). All the cell lines under study showed significantly reduced viability when treated with 5 µM G1. Notably, SW-620, which had the highest expression of GPER protein, showed significantly reduced viability in response to 1µM G1 (Fig 1B and Fig 3B). COLO-320DM, with no or negligible expression of GPER, did not show significant reduction of viability only at 1 µM G1 (Fig 1B and Fig 3E). The two cell lines significantly differed in their response to 5 µM G1. At this concentration the viability of SW620 was reduced by 8.44%, whereas the viability of COLO-320DM was reduced by 29.36% (Fig 3B and 3C).

**Figure 3.**
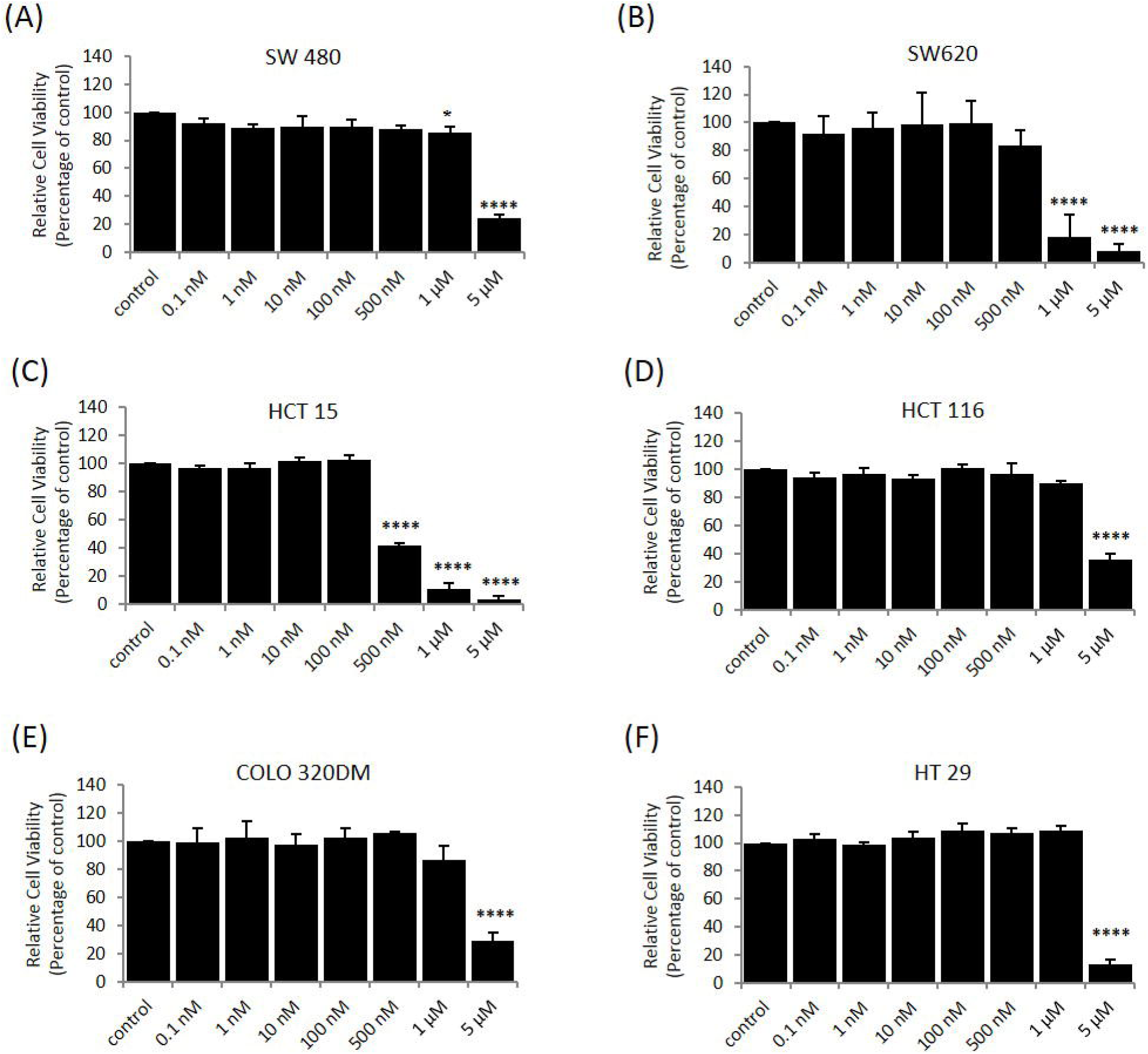
Effect of G1 treatment on viability of colon cancer cell lines. The indicated cell lines were treated with varying concentrations of G1 for a period of 120 h. The change in viability was determined as described in materials and methods. Bars represent mean relatively viability ± SD (n = 4 (SW620) and n = 3 (for other cell lines); each biological replicate comprising of 6 technical replicate wells for each of the treatment groups).

## DISCUSSION

The recognition of GPER as a mediator of estrogen effects not only brought complexity to estrogen signaling, but also ushered a new dimension to the etiology, progression, and therapeutic resistance of endocrine cancers. Recent studies have expanded the scope and relevance of GPER expression and regulation beyond the traditionally conceived endocrine (estrogen) responsive tissues (27). Two independent reports implicated GPER in the pathophysiology of colon cancer (31,32), although there is ambiguity about the role of GPER. The present work is set against the backdrop of growing literature on epigenetic silencing of GPER in diverse pathological contexts (31,33–35,40,41), and the proposed tumor suppressor role of GPER (31).

Liu and co-workers found that GPER down-modulation in colorectal cancer was associated with promoter methylation and histone deacetylation (31). They analyzed a CpG island (between −461 and −781) in the GPER promoter by bisulfite sequencing and found significantly greater proportion of methylated CpGs in colorectal cancer compared to normal tissues. This was corroborated by our independent analysis of the TCGA data (36). Liu and co-workers also analysed the CpG island in colorectal cancer cell lines. However, their conclusion was based on few sequences, with no statistical analysis. The region analyzed by Liu and co-workers overlapped with the upCpGi. However, it did not include the DMR, which we had described earlier in breast cancer cell lines (35,36). The technology used to generate TCGA methylation data does not have probes to interrogate the DMR. Hence, our analysis of TCGA colorectal cancer data was unable to provide information about methylation in the DMR. The present work was motivated by the need to address the following missing links-a) the status of methylation in the DMR in colorectal cancer cells, and b) the correlation between GPER expression and methylation in the upCpGi.

To our knowledge, this is the first study to analyze GPER expression in colon cancer cell lines and correlate it with the extent of methylation in the upCpGi, with special reference to the DMR. One of the striking observations from our bisulfite sequencing data is the concentration of methylated CpGs in the DMR of the upCpGi. This is reminiscent of the methylation pattern reported earlier in MCF-7 and MDA-MB-231 breast cancer cells. In the COLO-320DM cell line the methylated CpGs were found throughout the upCpGi. COLO-320DM and SW-620 cells represented two extreme states of methylation, and the remaining cell lines showed varying degrees of CpG methylation in the DMR. Interestingly, inserts in 6 clones from HCT-15 cells, which were sequenced showed the absence of the twenty-fifth CpG dinucleotide. Similarly, one clone from HT-29 had an absence of the first CpG dinucleotide (Fig. 2A). We believe that this is due to the heterogeneity brought about by mutations that accumulate during cell passages. The inverse correlation between methylation in the DMR and GPER expression suggests that DNA methylation dependent silencing of GPER could be the underlying cause of GPER downmodulation in colon cancer.

Estrogen’s oncoprotective effect on the colorectal tissue has a sound epidemiological basis. The receptors that transduce the protective effects of estrogen merit special attention on the following counts. First, they hold the key to our understanding of the effects of estrogen in a tissue traditionally considered non-endocrine. Second, they may inspire novel therapeutic strategies against colorectal cancer. Given the absence or undetectable expression of ERα in the cell lines used (data not shown), the study also attempted to correlate the status of methylation in the DMR or GPER expression with response to G1. We were not successful in finding a significant correlation. However, the pattern of methylation, GPER expression and response to G1 in COLO-320DM and SW-620 cells, which have contrasting levels of methylation and GPER expression, suggest that reduced GPER expression brought about by DNA methylation results in reduced response to GPER activation. GPER activation induces cell cycle arrest and apoptosis in colon cancer cells. Thus, we propose that DNA methylation dependent loss of GPER expression negatively impacts estrogen’s oncoprotective effect in the colonic epithelium, which contributes to colon cancer.

Tumor suppressors are frequently inactivated through methylation of DNA in neoplastic conditions. The biomarker and prognostic potential of methylated CpG islands in specific tumor suppressors are appreciated generally. Despite the ambiguity in the perceived role of GPER in the colorectal tissue, its expression level in primary tumors appears to be an important clinical marker. The present study confirms the inverse relationship between upCpGi methylation and GPER expression in cell line models. Furthermore, it underscores the importance of the DMR, which encompasses the terminal eight CpGs, in methylation-dependent silencing of GPER. More investigations are warranted to explore DMR hypermethylation as an indicator of GPER expression in colon cancer and its utility in diagnosis and prognosis.

## AUTHOR CONTRIBUTIONS

UP and AML designed the experiments. UP and SG performed the experiments. UP and AML analysed the results. UP, SG, and AML wrote the manuscript.

## ACKNOWLEDGEMENT

The work was supported by financial assistance from the Indian Council of Medical Research, Govt. of India (Sanction letter No. 2019-0687/CMB/ADHOC-BMS, dated 13.11.2019). Infrastructural support from the Department of Biosciences and Bioengineering at IIT Guwahati is acknowledged. SG was a student of the M.Tech program supported by the Department of Biotechnology, Govt. of India, which also provided her fellowship.

